# SLC26A3 (DRA) is stimulated in a synergistic, intracellular Ca^2+^ dependent manner by cAMP AND ATP in intestinal epithelial cells

**DOI:** 10.1101/2022.12.02.518884

**Authors:** Rafiquel Sarker, Ruxian Lin, Varsha Singh, Mark Donowitz, Chung Ming Tse

**Author notes:** Correspondence: C. Ming. Tse, PhD, Department of Medicine/GI Division, The Johns Hopkins University, Ross 930, 720 Rutland Street, Baltimore MD21205.

## Abstract

In polarized intestinal epithelial cells, DRA is a brush border (BB) Cl^-^/HCO_3_^-^ exchanger that is part of neutral NaCl absorption under baseline conditions but in cAMP driven diarrheas it is stimulated and contributes to increased anion secretion. To further understand regulation of DRA in conditions mimicking some diarrheal diseases, differentiated Caco-2/BBE cells were exposed to forskolin and ATP. Forskolin and ATP both acutely stimulated DRA in a concentration dependent manner. Forskolin at 1µM and ATP at 0.25 µM had minimal to no effect on DRA activity given individually; however, together, they stimulated DRA activity to levels seen with maximum concentrations of forskolin and ATP alone. In Caco-2/BBE cells expressing the Ca^2+^ indicator GCaMP6s, ATP alone increased Ca^2+^ in a concentration dependent manner, while forskolin (1 µM), that by itself did not significantly alter Ca^2+^, followed by 0.25 µM ATP produced a large increase in Ca^2+^ that was ∼equal to the elevation caused by 1 µM ATP. BAPTA-AM pretreatment prevented the ATP and forskolin/ATP synergistic increased DRA activity and the increase in intracellular Ca^2+^ caused by ATP/forskolin. Conclusion: In Caco-2/BBE cells subthreshold concentrations of forskolin (cAMP) and ATP (Ca^2+^) synergistically increased intracellular Ca^2+^ and stimulated DRA activity with both being blocked by BAPTA-AM pretreatment. Diarrheal diseases such as bile acid diarrhea, in which both cAMP and Ca^2+^ are elevated, are likely to be associated with stimulated DRA activity contributing to increased anion secretion, while separation of DRA from NHE3 contributes to reduced NaCl absorption.

## INTRODUCTION

SLC26A3 (DRA) is an intestinal brush border (BB) Cl^-^/HCO_3_^-^exchange protein that is linked to the BB Na^+^/H^+^ exchanger SLC9A3 (NHE3) to produce the neutral NaCl absorptive process, the major way intestinal Na^+^ absorption occurs in the period between meals (1-9). DRA is expressed in the largest amount in human proximal colon but is also present in significant amounts in the human duodenum and ileum (3,6,7, 10). It is localized to the upper crypt and villus in the small intestine and upper crypt and surface cells of the colon (3, 7,10); in these cells it is colocalized with NHE3 and in many of these cells, also with CFTR (3,12-15). DRA functionally and physically is linked to NHE3 and takes part in neutral NaCl absorption under baseline condition. However, in addition in some conditions associated with elevated cAMP, DRA also carries out increased rates of HCO_3_ ^-^ secretion and in the same cases physically interacts with the BB Cl channel, CFTR (3, 13) and dissociates from NHE3. However, the role of DRA in diarrheal diseases is unclear; DRA is inhibited in IBD and Salmonella related diarrheas, while its failure to function or failure to be produced causes congenital chloridorrhea (2, 5, 16,17)There is increasing evidence that increased second messengers, as occurs with most diarrheal diseases, alter DRA activity (2,3,5,8, 9,13,14,17-24). Elevation in intracellular cyclic nucleotides and/or Ca^2+^ are part of the pathophysiology of many diarrheal diseases; for instance, cAMP in cholera toxin related, cGMP in E. coli heat stable enterotoxin related, and Ca^2+^ in rotavirus related diarrhea, while in bile salt induced diarrhea both elevated cAMP and elevated Ca^2+^ contribute to the pathogenesis (25-31). In addition to the role of cAMP and elevated intracellular Ca^2+^ signaling pathways in pathologic processes, they also take part in regulation of multiple physiologic processes, in which their actions are independent, additive, synergistic or inhibitory. In the current studies, we tested the hypotheses that both elevated cAMP and Ca^2+^ stimulated DRA activity and examined the interactions of elevated cAMP and elevated Ca^2+^ in DRA stimulation to determine if they affected DRA entirely independently or if there were interactions that were additive or synergistic in a model polarized intestinal epithelia cell line.

## MATERIALS AND METHODS

### Chemicals, reagents, and antibodies

Reagents were obtained from Sigma-Aldrich (St. Louis, MO) unless otherwise stated. Ca^2+^ ionophore 4-Br-A23187 was from Biomol (Plymouth Meeting, PA). 2,7-Bis(2-carboxyethyl)-5-carboxyfluorescein acetoxymethyl ester (BCECF-AM) and Alexa Fluor 488 and 568–conjugated goat anti-mouse and antirabbit secondary antibodies were from Invitrogen (Carlsbad, CA). Mouse monoclonal antibodies to the HA epitope were from Covance Research Products (Princeton, NJ). Rabbit polyclonal antibodies to HA were from Santa Cruz Biotechnology (Dallas, TX). Mouse monoclonal FLAG antibody and Rabbit anti-NHERF2 antibodies were from Sigma-Aldrich. HA-cholinergic M3 receptor cDNA was from Addgene (# 40753). BPTU (1-(2-(2-(tert-butyl)phenoxy)pyridin-3-yl)-3-(4-(trifluoromethoxy) phenyl) urea was from Tocris Bioscience.

### Cell culture

The Caco-2/BBE cell line originally derived from a human colon adenocarcinoma was obtained from M. Mooseker (Yale University) and J. Turner (University of Chicago). Cells were grown on 0.4µ membrane Transwell inserts in DMEM containing 25 mM NaHCO_3_ supplemented with 0.1 mM nonessential amino acids, 10% fetal bovine serum (heat inactivated, 55°C for 30 min), 4 mM glutamine, 50 U/ml penicillin, and 50 μg/ml streptomycin, pH 7.4, in 5% CO_2_/air at 37°C.

### Stably transduced Caco-2/BBE cells with lentivirus GCaMP6s expression and transiently transduced with adenovirus-HA-NHE3

Caco-2/BBE wild-type cells were transduced with GCaMP6s lentiviral particles and a stable Caco-2/GCaMP6s stable cell line generated with puromycin selection (10 μg/mL) as described (32). Adenovirus-HA-NHE3 was prepared as described and studied ∼48 h after transduction (32)

### Measurement of Cl^-^/HCO_3_ ^-^ Exchange Activity

Cl^-^/HCO_3_^-^ exchange activity was measured fluorometrically using the pH_i_-sensitive dye BCECF-AM and a custom chamber allowing separate apical and basolateral perfusion, as previously described (14,33). Cells were incubated with 10 mmol/L BCECF-AM in NaCl solution (138 mmol/L NaCl, 5 mmol/L KCl, 2 mmol/L CaCl_2_, 1 mmol/L MgSO_4_, 1 mmol/L NaH_2_PO_4_, 10 mmol/L glucose, 20 mmol/L HEPES, pH 7.4) for 30 minutes at 37°C and mounted in a fluorometer (Photon Technology International, Birmingham, NJ). Cells were perfused on the apical surfaces with Cl^-^ solution (110 mmol/L NaCl, 5 mmol/L KCl, 1 mmol/L CaCl_2_, 1 mmol/L MgSO_4_, 10 mmol/L glucose, 25 mmol/L NaHCO_3_, 1 mmol/L amiloride, 5 mmol/L HEPES, 95% O_2_/5% CO_2_) or Cl^-^ -free solution (110 mmol/L Na-gluconate, 5 mmol/L K-gluconate, 5 mmol/L Ca-gluconate, 1 mmol/L Mg-gluconate, 10 mmol/L glucose, 25 mmol/L NaHCO_3_, 1 mmol/L amiloride, 5 mmol/L HEPES, 95% O_2_/5% CO_2_) at a flow rate of 1 mL/min; the basolateral perfusion contained the Cl^-^ solution throughout the experiment. The switch from Cl^-^ containing to Cl^-^ -free solution generates a Cl^-^ out concentration gradient which stimulates HCO_3_^-^ entry across the cell membrane performed by Cl^-^/HCO_3_ ^-^ exchanger(s), and the resulting rate of change in pH_i_ was recorded. Multiple rounds of removing (indicated by closed arrowheads) /replenishing (indicated by open arrowheads) extracellular Cl^-^ were performed to determine the Cl^-^/HCO_3_ ^-^ exchange activity under basal conditions as a time control as well as in the presence of several compounds, including forskolin (1-10 µM, apical and basolateral, 7min), ATP (0.25-10 µM, apical), UTP (1 µM, apical) and carbachol (1 µM, basolateral). At the end of each experiment, pH_i_ was calibrated using K-clamp solutions with 10 mmol/L nigericin (Cayman Chemical, Ann Arbor, MI) that were set at pH 6.8 and 7.8. The initial rate of alkalinization after the switch from Cl^-^ solutions to Cl^-^ - free solutions was calculated using Origin 8.0 software (OriginLab, Northampton, MA) and in some cases the stable pHi reached after alkalinization was determined. Apical Cl^-^/HCO_3_^-^ exchange in these Caco-2/BBE cells was shown to be primarily DRA based on prevention of the change in apical Cl^-^ removal-induced alkalinization by pretreatment with DRA inhibitor-A250 (10 µM).

### Intracellular Ca^2+^ measurements

Plasmid pGP-CMV-GCaMP6s (http://addgene.org/40753) was cloned into lentiviral vector pCDH-EF1-MCS-IRES (puro) and lentiviral particles produced. Caco-2/BBE wild-type cells were transduced with GCaMP6s lentiviral particles and stable Caco-2/GCaMP6s cell lines was generated with puromycin selection (10 μg/mL). Caco-2 cells stably expressing GCaMP were seeded into 12-well Transwell (polyester membrane, Corning 3460) plates (1 × 10^5^ cells/well). Cells grown for 12–15 days post-confluency were serum deprived for 2–3 h before study. Live images of GCaMP fluorescence were obtained every second with an Olympus FV30000RS confocal microscope with 10x/0.75NA objective lens using the resonance mode, at 37°C in the OkoLab stage top plus transparent shroud incubator. ATP (0.25 µM or 1 μM) was added apically at specific time points without interrupting the live cell imaging. Time series were acquired with 512 × 512 pixels, 300 × 300-μm field of view for 600 time points at ∼1 s per time point. In some studies, EGTA (2 μM) was added ∼1 min before forskolin and/or ATP addition. In other studies, cells were treated with BAPTA-AM (25 µM) for 20-30 min before forskolin and/or ATP addition. Intensity of GCaMP6_S_ fluorescence was quantitated by MetaMorph. To determine changes in Ca^2+^, ATP-induced changes in GCaMP6s fluorescence (*F*) were divided by baseline fluorescence (*F*_0_).

### Statistical Analyses

Results are expressed as means ± SEM. Statistical evaluation was by Student’s *t* tests or ANOVA with Bonferroni correction when three or more comparisons were made. *p* values ≤ 0.05 were considered significant.

## RESULTS

### DRA is acutely stimulated by elevated intracellular Ca^2+^

We previously demonstrated that cAMP (forskolin) acutely stimulated DRA activity in Caco-2/BBE cells and human colonoids (14). Here we determined the effect of elevating intracellular Ca^2+^ on DRA activity in these cells. Receptor mediated Ca^2+^ release from intracellular calcium stores, including endoplasmic reticulum, occurs through both intestinal P2Y purinergic receptors and cholinergic M3 receptors that includes activation of PLCs (35,36). Shown in Fig 1A/B, in which DRA was measured in the same Caco-2/BBE monolayer under basal conditions and after ATP (1µM) exposure on the apical surface, ATP rapidly stimulated DRA activity. 1µM ATP caused an ∼40% stimulation of basal BB DRA activity. Similar experiments determined the effects of 10uM and 0.25 µM ATP. The ATP induced stimulation at 10 µM ATP was slightly but not significantly greater than that at 1µM, while minimal (∼10%) ATP stimulation occurred with apical 0.25 µM ATP exposure (Fig 1C). The effect of apical addition of 1µM UTP was also determined. Similar to the effect of ATP, apical UTP rapidly stimulated DRA activity (Fig 1D). Because, rotavirus, which causes diarrhea and inhibits Na^+^ absorption as well as induces intestinal fluid secretion, was shown to cause calcium waves by activation of P2Y1 receptors (35), it was determined if pretreatment with the P2Y1 receptor antagonist BPTU prevented the ATP (1µM) stimulation of DRA activity. As shown in Fig 1E, BPTU (10µM) pretreatment did not alter ATP stimulation of DRA activity.

**Fig 1.**
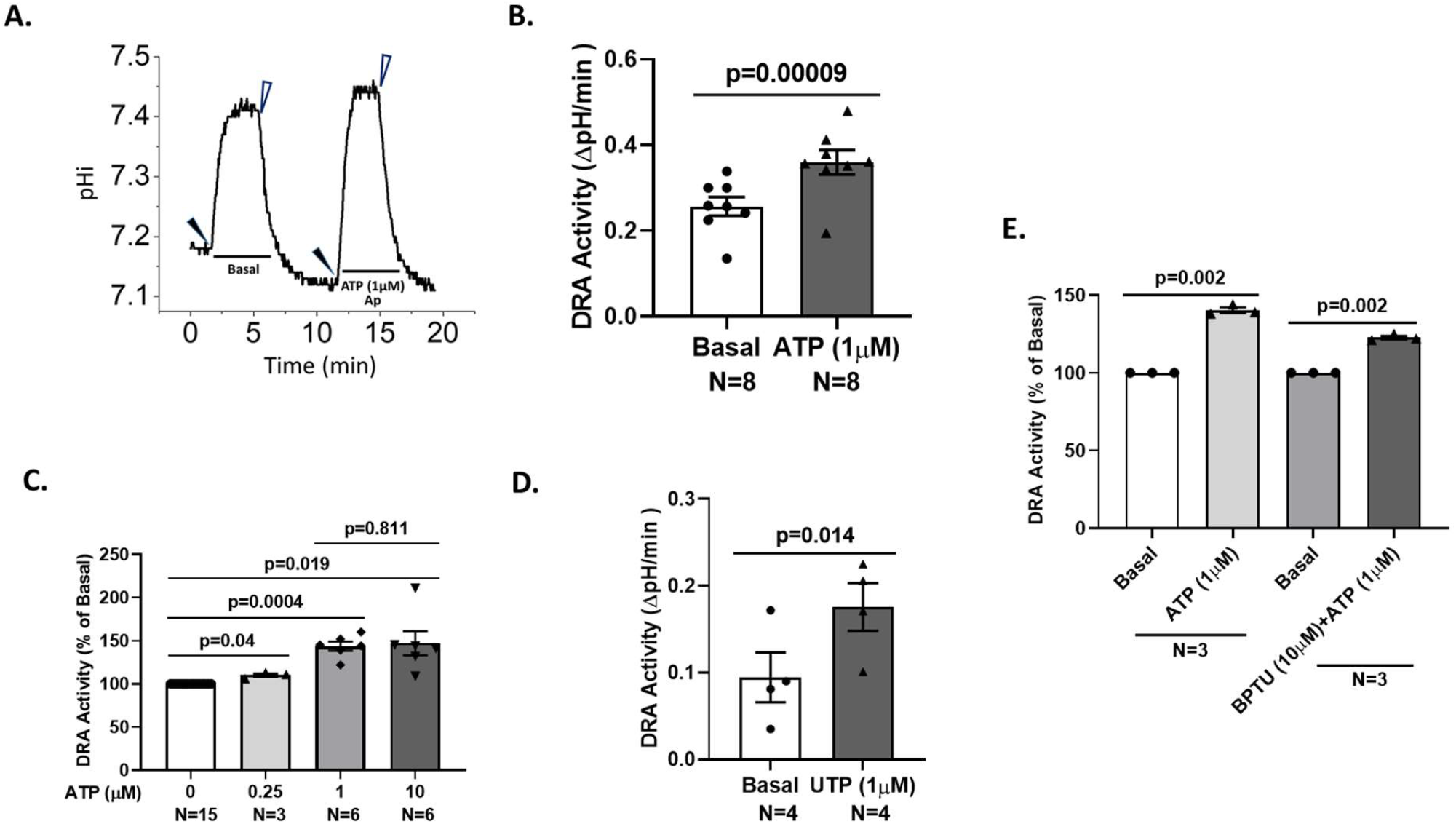
ATP and UTP acutely stimulate DRA activity in Caco-2/BBE cells. A. Cl^-^/HCO_3 -_ activity was determined by measuring intracellular pH_i_ with BCECF/fluorometry as apical bathing Cl^-^ solution removal stimulation of intracellular alkalinization that was reversed by re-addition of Cl^-^ after a constant pH_i_ was reached. Polarized Caco-2/BBE cells with apical and basolateral surface perfused with Cl^-^ reached a steady state pH_i_ and then Cl^-^ was removed only from the apical media and initial rates of intracellular alkalinization were determined and in some cases the steady state of pH_i_ reached was also determined. Multiple cycles of removal (closed arrowhead)/re-addition (open arrowhead) of Cl^-^ were performed and the effect of apical ATP (1µM) was determined. Results from a single monolayer are shown with increased initial rate and steady state pH_i_ after ATP. The Cl^-^/HCO_3 -_ exchange activity is considered to be due to DRA activity since Cl^-^ removal induced alkalinization in these cells was entirely prevented by treatment with DRA inhibitor A250 (10 µM) (14). B. Effect of ATP (1 µM) on Cl^-^/HCO_3 -_exchange activity, determined as in Fig 1.A. C. Experiments as in Fig 1A/B studying the effects of multiple concentrations of ATP which were compared to DRA activity in the same monolayer before ATP perfusion; results were calculated with untreated DRA activity set as 100% for each experiment. 0.25 µM ATP caused a minimal stimulation of DRA, while 1 and 10 µM ATP caused similar significant stimulation compared to basal activity. D. Apical UTP (1µM) acutely stimulated apical Cl^-^/HCO_3 -_ exchange activity. The magnitude of increased DRA activity (compare Fig 1B and 1D) caused by UTP was similar to that caused by ATP. E. Acute ATP stimulation of DRA activity was not prevented by pretreatment with 10 µM BPTU, a specific P2Y1 inhibitor. Results in B-E are means ±SEM. In all experiments, n represents the number of separate experiments.

Carbachol is known to elevate intracellular Ca^2+^ in intestinal epithelial cells via the M3 cholinergic receptor, acting on the ER through PLC-IP_3_ signaling (36). The effect of carbachol was determined on DRA activity in Caco-2/BBE cells stably transduced to express the cholinergic M3 receptor, which is not present endogenously (Fig 2A). Addition of 1µM carbachol to the basolateral surface rapidly stimulated DRA activity (Fig 2B/C) by ∼45% above basal activity in the same cells. These results demonstrate that receptor mediated intracellular elevation of Ca^2+^ stimulates DRA activity in polarized human intestinal epithelial cells.

**Fig 2.**
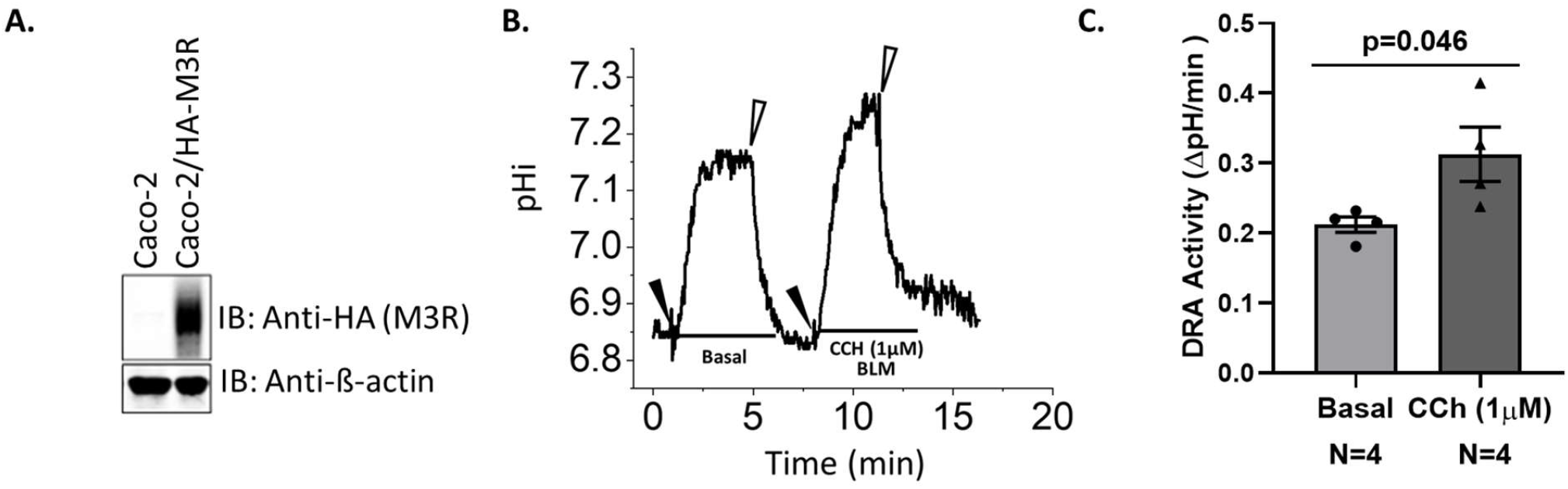
Carbachol acutely stimulates DRA activity in Caco-2/BBE-M3 receptor cells. A. Caco-2 cells were stably transduced with the cholinergic M3 receptor epitope tagged with HA, which was lacking endogenously. Lysates from these cells were immunoblotted for the M3 receptor, along with the loading control, β-actin. B. C. Experiments similar to those described in Fig 1A were performed but examining the effect of basolateral carbachol on Cl^-^/HCO_3 -_exchange activity in Caco-2/BBE-M3 cholinergic receptor cells. B. Single experiment; C. Means ±SEM of multiple experiments, comparing basal and carbachol effects on Cl^-^/HCO_3_^-^ activity from the same monolayers. n represents the number of separate experiments.

Having shown that acutely elevated Ca^2+^ stimulates DRA activity, it was determined if there was an interaction in the DRA stimulation by elevated cAMP and elevated Ca^2+^; specifically, whether Ca^2+^ elevation and increased cAMP affected DRA entirely independently or if there were interactions such that their effects were additive, synergistic, or competitive.

### cAMP and ATP acutely stimulate DRA in a synergistic manner

We previously reported that 10 µM forskolin acutely stimulated DRA in Caco-2 cells and in human colonoids (14) and confirmed that finding (Fig 3A/B). To evaluate the interactions of cAMP and elevated Ca^2+^ in the acute stimulation of DRA, lower concentrations of forskolin were evaluated for effects on DRA activity in comparison to the effects of the higher concentration. Fig 3C demonstrates that in comparison to DRA stimulation by 10 µM forskolin, exposure to 1 µM did not have a significant effect on DRA activity.

**Fig 3.**
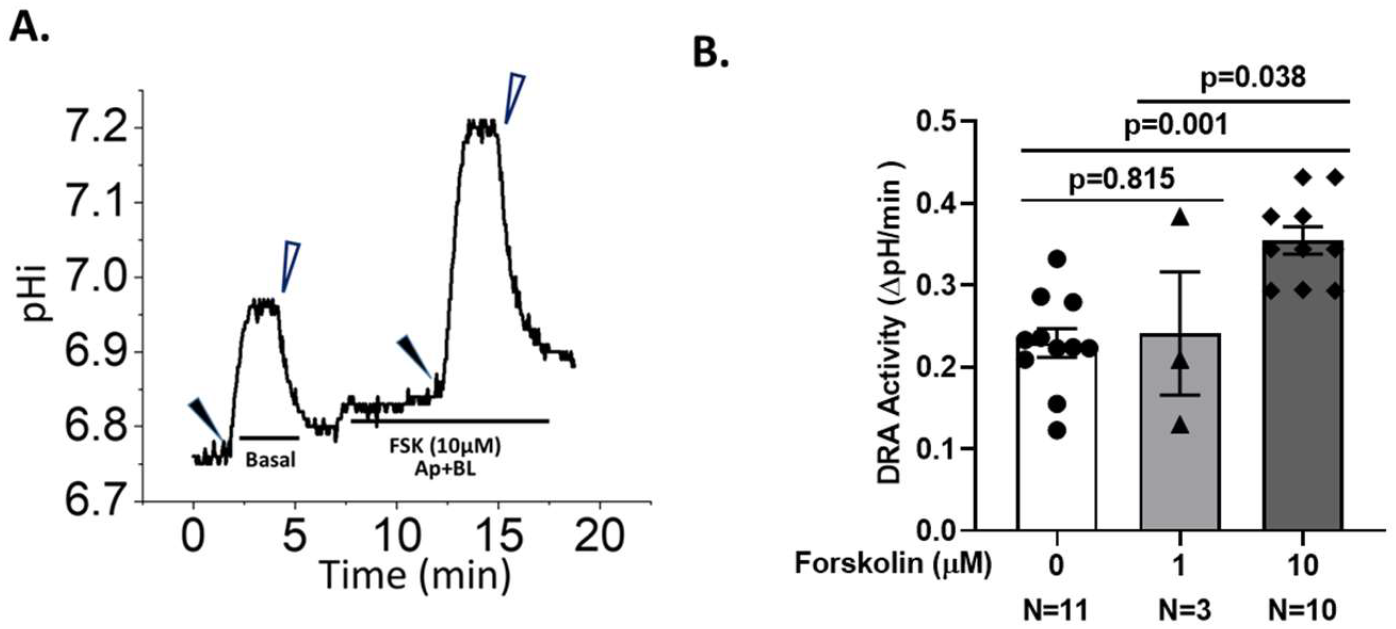
Acute DRA stimulation by forskolin in Caco-2/BBE cells. Apical and basolateral forskolin (10 µM) was exposed to Caco-2 cells and ∼7 min later effects on DRA activity were determined as in Fig 1A by apical Cl^-^ removal. A shows the Cl^-^/HCO_3 -_ activity in a single experiment. B. DRA activity, means ±SEM from multiple experiments comparing the effects on DRA activity of 1 and 10 µM forskolin, demonstrating significant stimulation of DRA by 10 µM but not 1 µM forskolin.

SLC26A6 (PAT-1) and CFTR were previously shown to be stimulated in a synergistic manner by elevation of cAMP and Ca^2+^ (13,37). Since 0.25 µM ATP caused a very minimal stimulation of DRA, while 1 µM forskolin did not alter DRA activity, the additive effects at these concentrations were determined. In these studies, forskolin (1µM) was exposed to the Caco-2/BBE cells for ∼7 min in Cl^-^ containing media and then ATP (0.25 µM) was added in a Cl^- -^free solution to determine DRA activity. While 1µM forskolin did not alter DRA activity, this combination of forskolin and ATP significantly increased DRA activity by ∼70% above baseline (Fig 4 A-C). These results demonstrate that concentrations of forskolin and ATP which themselves cause minimal or no effects on DRA activity, together cause a robust stimulation, thus demonstrating a synergistic interaction of cAMP and Ca^2+^. Further studies examined the effects of maximal concentrations of forskolin (10µM) and ATP (10 µM) alone and together on DRA activity. While both 10 µM ATP (140 ± 2%) and 10 µM forskolin (164±6%) alone acutely stimulated DRA, added sequentially, as done in Fig 4A, their DRA stimulation was not additive (184±11%) (Fig 4D).

**Fig 4.**
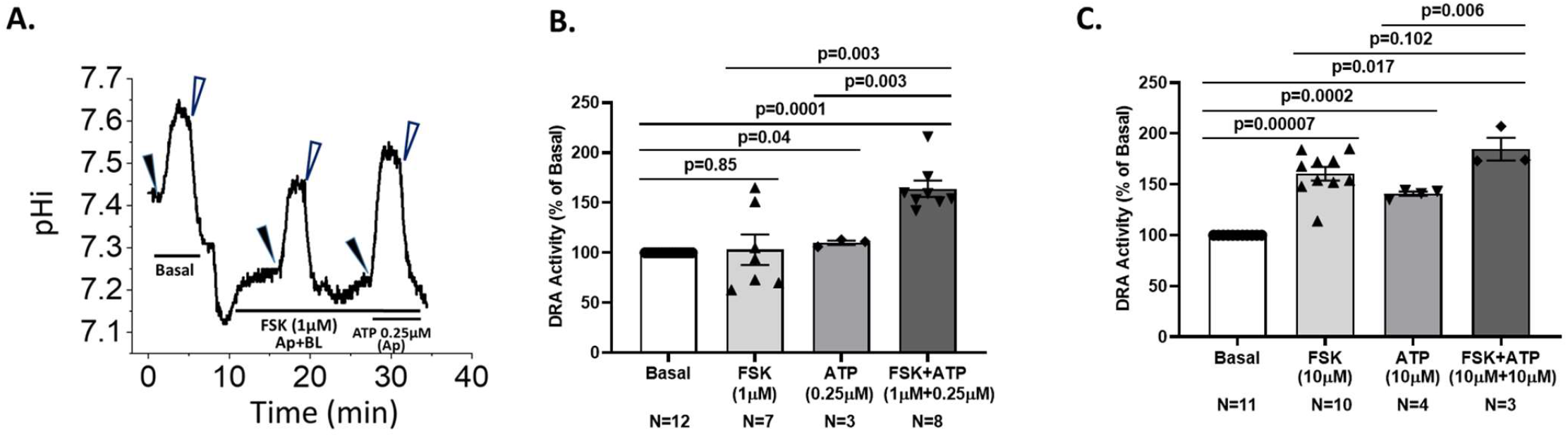
Concentrations of forskolin and ATP that have minimal effects on DRA activity cause synergistic stimulation of DRA in Caco-2/BBE cells. A. Single experiment in which Cl^-^/HCO_3 -_ exchange activity was determined under basal, untreated conditions and after forskolin (1 µM) addition to the apical and basolateral perfusions followed in ∼7 min by apical addition of ATP (0.25 µM). B. Multiple experiments as in A, showing DRA activity as means±SEM. n is the number of separate experiments. While neither forskolin (1µM) nor ATP (0.25µM) alone significantly altered DRA activity, together DRA activity was stimulated. C. Multiple experiments as in A and B showing DRA activity as means±SEM but with concentrations of forskolin (10 µM) and ATP (10 µM) that each maximally stimulate DRA activity and also studied together with a time course as in Fig 4A. Maximum concentrations of DRA plus ATP did not demonstrate additive or synergistic DRA stimulation. Results are means±SEM. n is the number of separate experiments.

Since DRA has been shown to take part in both neutral NaCl absorption, interacting with NHE3, and anion secretion, while the Caco-2/BBE cells used in these studies express a minimal amount of endogenous NHE3, we determined whether increased NHE3 expression altered the forskolin/ATP synergistic DRA stimulation. Adenoviral-NHE3 transduction was used to transiently increase NHE3 expression. The trace of a representative experiment in Fig 5A demonstrates that DRA stimulation in NHE3 expressing Caco-2/BBE cells with ATP (1µM) is characterized both by an increase in initial rate of DRA activity and an increase in the steady state pH_i_ reached and this is similar to wild type cells (Fig. 1A). Stimulation of DRA by forskolin and ATP using concentrations that produced synergistic stimulation, as described in Fig 4, also caused synergistic stimulation of DRA activity which was slightly but not significantly smaller than that which occurred without the increase in NHE3 expression (compare Fig 5B, 49 ±5% and Fig 4B, 64±8%, ns).

**Fig 5.**
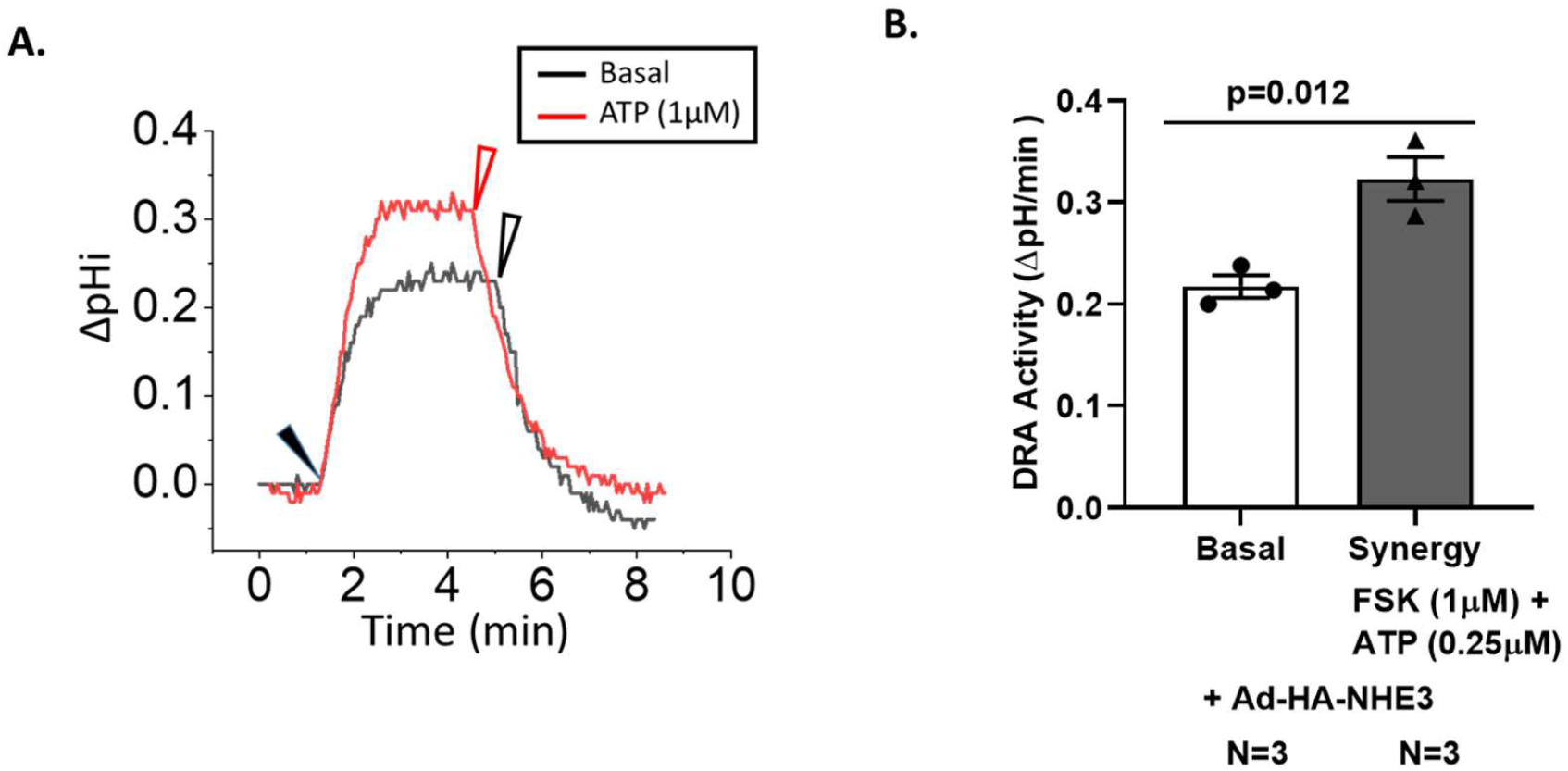
Forskolin/ATP mediated synergistic stimulation of DRA is similar in Caco-2/BBE cells acutely transduced with HA-NHE3. Experiments were performed to determine if the presence of HA-NHE3 in Caco-2/BBE cells affected the forskolin/ATP induced synergistic stimulation of DRA. Caco-2 cells was transduced with adenovirus-HA-NHE3 and studied for Cl^-^/HCO_3_^-^ exchange activity ∼48 h later. A. Single experiment from the same Caco-2 monolayer with DRA activity determined under basal conditions and after apical addition of 1µM ATP. The traces are superimposed starting at time 0, although they were performed sequentially. Note that ATP stimulated the initial rate of DRA as well as increased the steady state pH_i_ achieved. B. Similar studies as those performed in Fig 4 with basal DRA activity, effects of adding forskolin (1 µM) followed by ATP (0.25 µM) demonstrated. Results are means±SEM. n is the number of separate experiments.

### Cytosolic free Ca^2+^ is increased by ATP alone and synergistically increased by the low concentrations of forskolin and ATP which synergistically stimulate DRA

Changes in free intracellular Ca^2+^ caused by ATP alone and following forskolin were determined in Caco-2/BBE cells stably expressing the genetically encoded Ca^2+^ indicator, GCaMP6s, that responds to elevation in free Ca^2+^ with an increase in fluorescent intensity. GCaMP6s was stably expressed in Caco-2/BBE cells by lentiviral transduction with puromycin selection which created a cell line with GCaMP6s expression in nearly all cells, as demonstrated by confocal microscopy (Fig 6A). ATP, when added apically to these cells, caused a very rapid increase in intracellular Ca^2+^ that occurred in a concentration dependent manner between 0.25 µM and 10 µM ATP. 0.25 µM ATP caused a single small peak in Ca^2+^ elevation that occurred in some cells and even this partial response only occurred in some but not all experiments. One and 10 µM ATP caused higher initial peaks in Ca^2+^ but also induced several small Ca^2+^ peaks that occurred at later times (Fig 6B). In contrast to the effect of ATP, forskolin at 1 (Fig 7) and 10 µM (data not shown) did not alter Ca^2+.^

**Fig 6.**
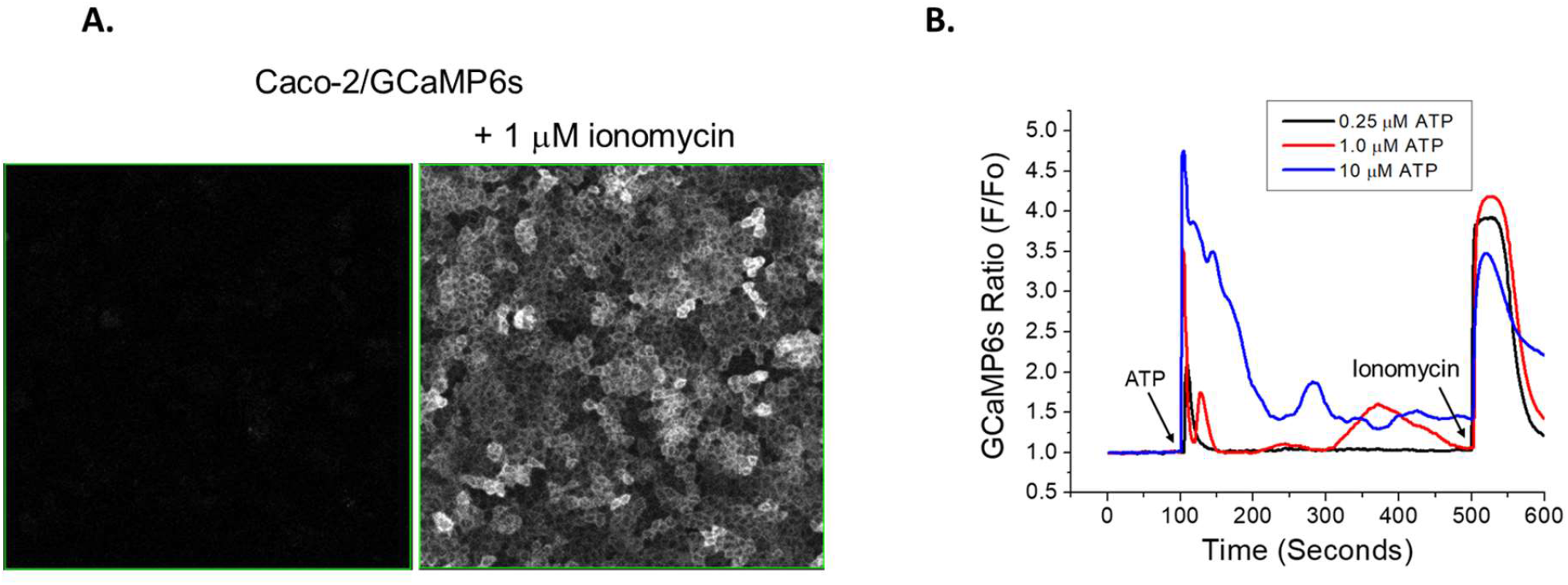
Apical ATP causes a concentration dependent increase in intracellular Ca^2+^ in Caco-2/BBE cells stably expressing the Ca^2+^ indicator GCaMP6s. A. The Ca^2+^ sensor protein GCaMP6s was stably expressed in Caco-2/BBE cells grown on glass bottom dishes and the effects of several concentrations of apical ATP was determined on intracellular Ca^2+^ in cells in an OkoLab stage top plus transparent shroud incubator, studied with Olympus FV30000RS confocal microscope (20x/0.75na objective) using the resonance mode at 37°C ; exposure to ionomycin (1 µM) produced an intracellular fluorescence signal demonstrating nearly uniform expression of GCamP6s in these cells. B. Exposure to several concentrations of apical ATP was quantitated as the ratio of the maximum fluorescence signal achieved to that originally present (F/Fo); ionomycin (1µM) was added as a positive control at the end of study of each monolayer. ATP caused a concentration dependent increase in intracellular Ca^2+.^ Single traces of each concentration of ATP are shown.

**Fig 7.**
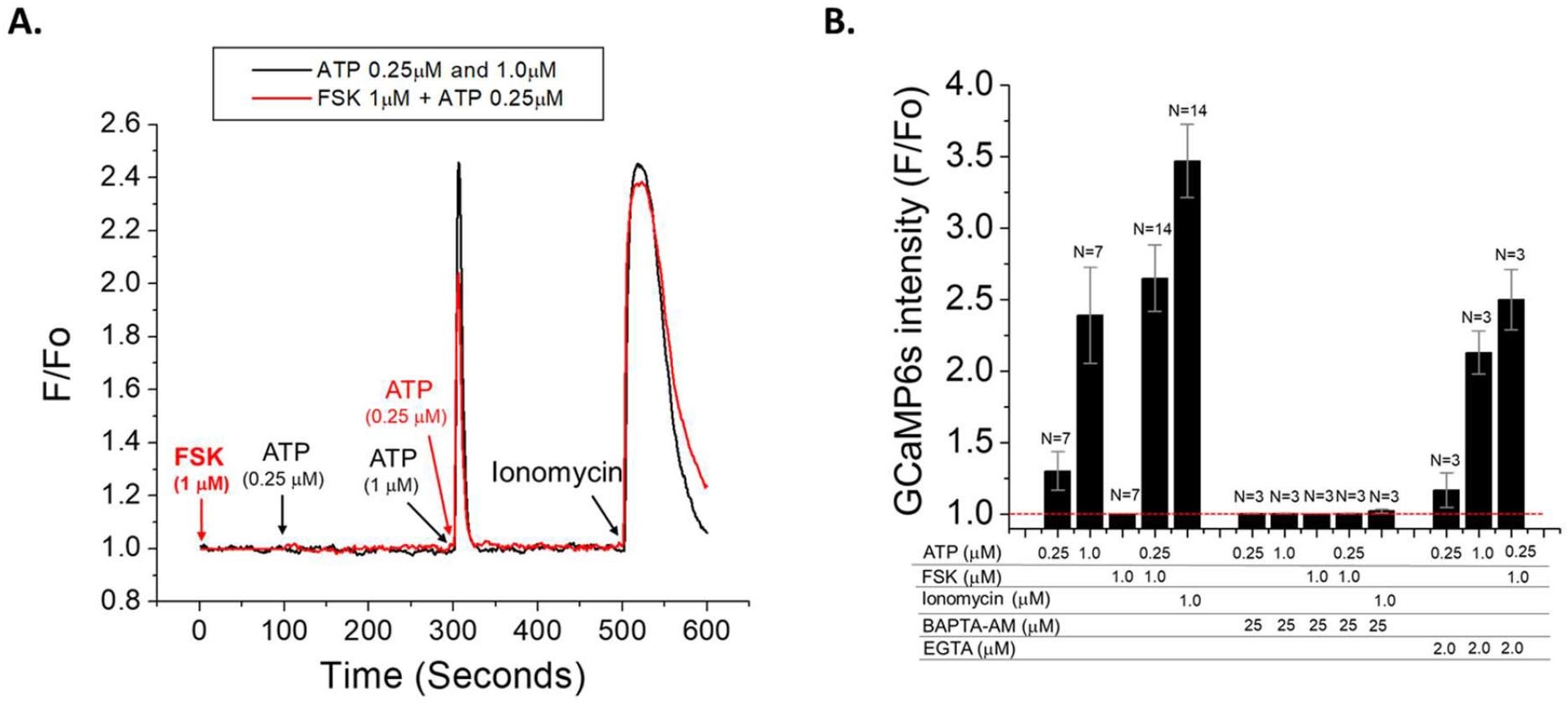
Forskolin and ATP synergistically increase intracellular Ca^2+^ in Caco-2/BBE-GCaMP6s cells. A. Intracellular Ca^2+^was determined as in Fig 6 in Caco-2/BBE-GCaMP6s cells exposed to 0.25 µM ATP followed by 1 µM ATP and then after 1 µM ionomycin as a positive control. In parallel, the forskolin (1 µM)/ATP (0.25 µM) conditions that led to synergistic stimulation of DRA were studied. Synergistic forskolin/ATP produced elevated Ca^2+^ similar to that produced by 1 µM ATP and that produced by ionomycin. B. Intracellular Ca^2+^ in Caco-2/BBE-GCaMP6s cells showing maximum increased Ca^2+^ caused by ATP (0.25 and 1 µM), forskolin (1 µM), and synergistic concentrations of forskolin (1 µM)/ATP (0.25 µM); studies also performed after pretreatment with BAPTA-AM (25 µM, 20 -30 min) or EGTA (2 µM, 1 min) added before ATP and/or forskolin. Ionomycin (1 µM) was added as a positive control. Results are means ±SEM of maximum Ca^2+^ level. n is the number of monolayers studied. BAPTA-AM prevented any changes in intracellular Ca^2+,^ while EGTA pretreatment did not alter the increase in Ca^2+^ caused by 1 µM ATP and by forskolin (1 µM)/ ATP (0.25 µM).

Whether the synergistic stimulation of DRA by forskolin and ATP was reflected by a synergistic increase in intracellular Ca^2+^ was determined. Addition of forskolin (1 µM) and ATP (0.25 µM) with the same time sequence as in Fig 4 caused a rapid increase in intracellular Ca^2+^ following ATP addition (Fig 7A). This increase was similar in magnitude to the effect of 1 µM ATP and to the increase caused by 1µM ionomycin, that was used as a positive control (Fig7 A, B). The elevation of intracellular Ca^2+^ by ATP (0.25 and 1 µM) and the forskolin/ATP concentrations causing the synergistic increase in Ca^2+^ were totally prevented by pretreatment of the Caco-2/BBE cells with BAPTA-AM (25 µM, 20-30 min pretreatment) (Fig 7B), while exposure to 2µM EGTA for 1 minute before ATP exposure did not alter the 1µM ATP-induced increase in Ca^2+^ or the synergistic increase caused by forskolin (1 µM)/ATP(0.25 µM) (Fig 7B).

### Elevated Ca^2+^ is necessary for synergistic stimulation of DRA by forskolin/ATP

Whether the synergistic stimulation of DRA activity was Ca^2+^ dependent was determined, using the same conditions that were demonstrated to prevent the forskolin/ATP increase in intracellular Ca^2+^. Pretreatment of Caco-2/BBE cells with 25 µM BAPTA-AM for 20-30 min abolished the synergistic stimulation of forskolin (1µM)/ATP (0.25 µM). In addition, BAPTA-AM pretreatment decreased basal DRA activity, although there was a wide range of basal DRA activities in different experiments. (Fig 8)

**Fig 8.**
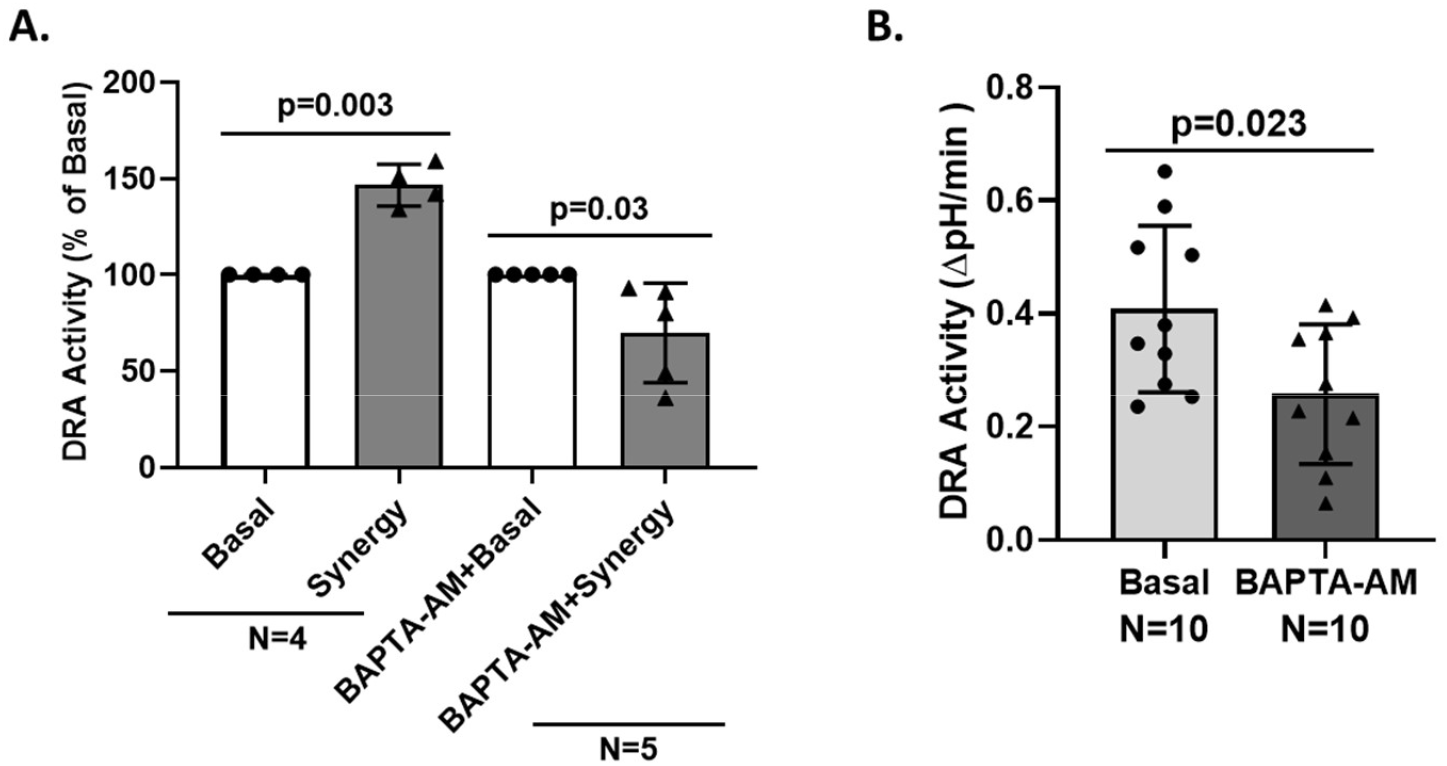
BAPTA-AM prevents the forskolin/ATP synergistic increase in DRA activity. **A**. Cl^-^/HCO_3_^-^ exchange activity was determined in Caco-2/BBE cells under basal conditions and when stimulated by forskolin (1 µM)/ATP (0.25 µM) as in Fig 4. Similar studies were performed in monolayers exposed to BAPTA-AM (25 µM, 20-30 min pretreatment). BAPTA-AM pretreatment prevented the synergistic stimulation of DRA. Results are means ±SEM. n is the number of separate experiments. B. Comparison of basal DRA activity from experiments including those in A, demonstrating that BAPTA-AM significantly reduced basal DRA activity.

## DISCUSSION

This study demonstrates that the forskolin and apical ATP induced elevated Ca^2+^ acts synergistically to acutely stimulate DRA in polarized intestinal epithelial cells. Synergistic interaction of cyclic nucleotide and Ca^2+^ signaling is not unique for regulation of DRA in intestinal epithelial cells. In fact, cAMP/Ca^2+^ synergy occurs in many aspects of signal transduction across species, organs, and physiologic responses as well as is involved in transcriptional and post-transcriptional signaling. Several examples include, effects on odor detection in drosophila (38); synergistic GH release from chicken pituitary (39); carbaprostacyclin (cPG12) induction of differentiation of preadipocytes to mature adipocytes (40); stimulation of c-FOS in Jurkat cells (41); and PGE2 -induced stimulation of prolactin transcription in Jurkat cells (42). While cAMP/Ca^2+^ synergy occurs in many signaling pathways, the cross-talk between the two second messenger systems does not always involve synergy; for example cAMP and Ca^2+^ are additive but not synergistic in ERK activation in a mouse model of long lasting long term potentiation, related to learning (43), and there are examples in which cAMP and elevated Ca^2+^ have opposing effects on the same signaling pathway, such as in metabotropic glutamate receptor signaling (44).

Intracellular Ca^2+^ was necessary for the cAMP/Ca^2+^ synergistic acute stimulation of DRA since both the effect of ATP alone on DRA and the synergistic stimulation were inhibited by pretreatment with BAPTA-AM but not by extracellular EGTA at a concentration that greatly lowered the extracellular free Ca^2+^ but was only present for a minute before ATP addition so as not to lower intracellular Ca^2+^. While intracellular Ca^2+^ was necessary for the synergy, we have not demonstrated that is sufficient and our studies have not considered additional factors that might be required for synergistic DRA stimulation, particularly those downstream from the Ca^2+^ elevation. Given the similar effect of ATP and UTP in stimulating DRA activity, dependence on intracellular and not extracellular Ca^2+^, and failure of the P2Y1 antagonist BPTU to inhibit the ATP induced increase in DRA activity, we conclude that other apical P2Y receptors are involved in the ATP elevation of Ca^2+^. Caco-2 cells, in addition to expressing low amounts of mRNA for P2Y1 receptors (10), with ADP being its preferred ligand, also express P2Y2, P2Y4, P2Y6 receptors (45,46). The three latter P2Y receptors all interact with Gq and activate phospholipase C isoforms, with ATP and UTP being their preferred ligands. Since UTP predominantly binds to P2Y2 and P2Y4 receptors, and to a lesser extent to P2Y6 receptors for which the preferential agonist is UDP, it is mostly likely that P2Y2 and/or P2Y4 receptors are responsible for ATP stimulation of DRA in Caco-2/BBE cells.

Multiple mechanisms for crosstalk/synergy between cAMP/Ca^2+^ signaling have been previously defined in detail. Given the involvement of apical P2Y receptors with PLC-IP_3_ generation, the studies of Muallem (47) are most relevant to our demonstration that synergistic stimulation of DRA requires the magnitude of elevation of intracellular Ca^2+^ to also increase synergistically. In the Muallem studies, cAMP increased the binding and sensitivity of IP_3_ to its ER Ca^2+^ channel receptor, IP_3_R, increasing the amount of Ca^2+^ released per molecule of bound IP_3_, to synergistically elevate cytosolic Ca^2+^. However, multiple other mechanisms of cAMP/Ca^2+^ synergy have been identified. These include, among others, Ca^2+^ dependent adenylyl cyclase activity (48); and direct effects of calmodulin binding to the CFTR R domain, with Ca^2+^-calmodulin binding increasing the open probability similarly to effects of cAMP-induced stimulation via phosphorylation, with overall the effects of Ca^2+^/CaM and cAMP being additive but together achieving the maximum activity caused by either alone (49). The latter results indicate effects via a common pathway. We have previously reported that FSK/cAMP elevates intracellular Ca^2+^via Epac1 in T84 cells (50). However, in the present study, we showed that FSK failed to increase intracellular Ca^2+^ (Fig.7A).

The current studies appear to help explain previous studies in the model cell, Caco-2/BBE, that was used in these studies, in which DRA seemed to be both stimulated and inhibited by cAMP (8, 14,15). We previously reported, based on use of proximity ligation assays in Caco-2/BBE cells expressing transduced NHE3 and endogenous DRA and CFTR, that under baseline conditions DRA bound NHE3 and CFTR and NHE3 bound DRA and CFTR; with elevation of cAMP the interaction of NHE3 with DRA and CFTR was reduced and that between DRA and CFTR was increased, with overall activity of DRA increasing (3). The seeming contradiction of both DRA stimulation and inhibition can be explained by occurrence of reduced neutral NaCl absorption related to the decrease in NHE3 on the plasma membrane reducing a pool that interacts with BB DRA and consequently reducing neutral NaCl absorption as demonstrated in Caco-2 cells in which Musch et al (8) used isotopic studies with ^22^Na and ^36^Cl to measure neutral NaCl absorption. Our studies indicate that in Caco-2/BBE cells the elevation of cAMP leads to increased DRA trafficking to the apical membrane where DRA interacts with CFTR and leads to increased anion secretion, as previously shown by us (14). Assuming that this occurs in cAMP/Ca^2+^ synergy in normal intestinal epithelial cells as well as in Caco-2 cells, it will be important to define the magnitude of the separation of NHE3 from DRA (which is part of the mechanism of inhibition of intestinal Na absorption that occurs in diarrhea as NHE3 trafficking from the plasma membrane to the early endosomes) and that of increased binding of DRA with CFTR (which is part of the stimulation of anion secretion). In the present studies, we showed that synergistic stimulation of DRA was observed whether NHE3 was present or not, and interestingly the magnitude of synergistic stimulation in the presence of NHE3 appeared to be slightly less.

The results presented here differ from the detailed studies of Lamprecht also using Caco-2/BBE cells (24,51). His studies acutely elevated intracellular Ca^2+^ with ionomycin or UTP and found that both did not alter initial rates of DRA activity but lowered the steady state pH_i_ achieved after ionomycin/UTP. This was interpreted to indicate that elevated intracellular Ca^2+^ probably changed the pH_i_ sensitivity of DRA as the major way Ca^2+^ altered DRA activity. What explains the differences in our findings that both forskolin and apical ATP as well as synergistic concentrations of forskolin and ATP stimulate DRA activity can only be speculated about. The current studies, however, show DRA stimulation that involves both initial rates and steady state pH_i_; effects that were reversed by BAPTA-AM, demonstrating that potential injury from elevated Ca^2+^ was not occurring. It must be noted that another major difference in these two Caco-2/BBE studies relates to the mechanism of DRA stimulation. In the Lamprecht study (24), neither ionomycin nor UTP altered the plasma membrane expression of DRA, while we previously reported that, using super-resolution combined with confocal microscopy, changes in surface expression of DRA and NHE3 supported that synergist stimulation of DRA was associated with increased apical membrane DRA expression and reduced NHE3 surface expression (3). Of note there was a major difference in the ways our studies were performed, consisting of the support the Caco-2 cells were grown on. In our studies, Caco-2 cells were grown on semipermeable supports (Transwell filters) allowing access of both apical and basolateral surface to bathing solutions with standardized polarization based on days post-confluency studied. Previous demonstration of Caco-2 polarization and differentiation has generally been done with cells grown on such semipermeable supports rather than the solid supports apparently used in the Lamprecht studies (24,51).

The demonstration that cAMP and Ca^2+^ synergistically stimulate DRA activity has relevance to pathophysiologic mechanism of multiple diarrheal diseases in which both cAMP and Ca^2+^ are elevated. In each of these there is a potential role for cAMP/Ca^2+^ cross-talk/synergy but in none have detailed evaluation of these interactions been achieved. PGE2 acutely elevates both intracellular Ca^2+^ and cAMP in intestinal epithelial cells and is part of the pathophysiology of IBD (52,53), with effects on multiple intestinal cell types that likely contribute to the IBD pathophysiology (ENS, macrophages, dendritic cells, others). Bile salts signal by cAMP and PKC as part of cholerrheic enteropathy in which increased bile salt concentration in the lumen of the colon stimulates CFTR activity while inhibiting NaCl absorption (54.55). In mouse B cells, cross linking the B cell antigen receptor using anti-IgM antibody initially elevates cAMP and subsequently increases intracellular Ca^2+^ leading to synergistic increase in CD80 transcription, which is part of B cell activation/interferon gamma production, that participates in multiple viral diarrheas, including rotavirus (56). In addition, low levels of apical ATP are known to be present in some polarized cells (57,58) with the cause being less certain although potential explanations include cell turnover/death, due to translocation of immunologic and inflammatory cells through the tight junctions, effects of interactions with the microbiome, among others. These low levels of ATP potentially are able to act synergistically with changes in cAMP that occur as part of digestion, as well as in diarrheal diseases, to synergistically participate in what is considered normal physiologic regulation of intestinal salt and water transport. Future studies evaluating the relevance of increases in both cAMP and Ca^2+^ in the same intestinal cells or in the multiple intestinal cells beyond the enterocytes that contribute to diarrheal disease pathophysiology are likely to provide new understanding of the pathophysiology of diarrhea and to provide drug targets to develop for treating diarrheal diseases.

## Abbreviations

BB: brush border;
IP3: Inositol 1,4,5-trisphosphate;
UTP: uridine 5’-triphosphate;
UDP: uridine diphosphate;
ATP: adenosine 5’-triphosphate;
PLC: phospholipase C;
CaM: calmodulin;
PGE2: prostaglandin E2;
IBD: inflammatory bowel disease;
BPTU: N-[2-[2-(1,1-Dimethylethyl)phenoxy]-3- pyridinyl]-*N*’-[4-(trifluoromethoxy)phenyl]urea;
ENS: enteric nervous system;
BAPTA-AM: 1,2-Bis(2- aminophenoxy)ethane-N,N,N’,N’-tetraacetic acid tetrakis(acetoxymethyl ester);
DRA: down-regulated in adenoma;
NHE3: Na+/H+ exchanger isoform-3

## ACKNOWLEDGEMENTS

Studies were supported in part by NIH/NIDDK RO1 DK26523, RO1 DK116352, R24 DK099803, PO1AI125181, and Digestive Disease Research Core Center Grant P30 DK089502. We acknowledge the helpful input of Shmuel Muallem, Ph.D in designing and interpretation these studies.

